# Voluntary alcohol consumption disrupts coupling of prefrontal cortical activity to arousal

**DOI:** 10.1101/2021.02.01.429229

**Authors:** Grayson Sipe, Ivan Linares-Garcia, My Nguyen, Elena Vazey, Rafiq Huda

## Abstract

Alcohol use disorder (AUD) exacts a major personal, societal, and economic toll. Top-down control from the prefrontal cortex (PFC), a critical hub for decision making, executive, and other cognitive functions, is key for the regulation of alcohol consumption. Arousal exerts profound effects on cortical processing, allowing it to potentially modulate PFC functions relevant for alcohol consumption and AUD. Despite this, it is unclear whether and how arousal-mediated modulation of PFC circuits relates to voluntary alcohol drinking behaviors. Two-photon microscopy is ideally suited for dissecting the neural circuit mechanisms underlying the effect of alcohol on intact circuits in behaving animals. We addressed a major limitation of this technology by developing a novel behavioral paradigm for voluntary drinking in head-fixed mice. We recorded responses of layer 2/3 excitatory neurons in the anterior cingulate cortex (ACC) subdivision of the PFC as mice voluntarily consumed ethanol, along with video recording of the pupil to track momentary fluctuations in arousal. Ethanol consumption bidirectionally modified the activity of subsets of ACC neurons, both at slow (minutes) and fast (sub-second) time scales. Remarkably, we found that the coupling of arousal to ACC activity before drinking was associated with subsequent ethanol engagement behavior. In turn, ethanol consumption modulated neuronal-arousal coupling. Together, our results suggest neuronal-arousal coupling as a key biomarker for alcohol drinking and lays the groundwork for future studies to dissect the therapeutic potential of this process for AUD and other substance use disorders.

## Introduction

Alcohol use disorder (AUD) is a debilitating set of chronic behaviors related to alcohol dependence that lead to long-term health, social, and economic detriments. Behavioral paradigms in which animals voluntarily self-administer ethanol are critical for understanding the molecular, cellular, circuit, and systems level neuronal processes underlying alcohol-related behaviors. Binge alcohol drinking is a pattern of alcohol consumption that can lead to the development of AUDs, however the mechanisms that predispose binge drinkers to develop AUDs are poorly understood. Although a wealth of mechanistic information has been generated by the various volitional drinking paradigms^1-6^, little is known about how *in vivo* neurophysiology actively changes while an animal drinks alcohol.

The prefrontal cortex (PFC) has canonical roles in decision making, attention, reward/punishment processing, social behavior, and other functions^7-9^. Consistent with the association of AUD with the dysfunction of these higher-order processes^10-13^, the PFC is highly implicated in AUD in humans^14^ as well as in non-human primate^15^ and rodent models^16-18^. Significant evidence describes mechanistic changes in PFC structure and function following acute or chronic ethanol exposure^19,20^, with its effect usually assessed in separate cohorts of animals at various time points throughout the exposure paradigm. Understanding how neuronal processing in PFC networks on the sub-second time scale relevant for ethanol consumption behavior relates to voluntary drinking and tracking this relationship across drinking is crucial for identifying the key neuroadaptations that contribute to the development of AUD.

Many PFC-driven behaviors are modulated by internal states, including by momentary fluctuations in arousal that can be approximated via changes in pupil size^21-25^. In turn, cortical activity is highly influenced by changes in arousal, with pupil dilations correlated to both increased and decreased activity relative to periods of pupil constrictions^24-29^. The coupling of cortical activity to arousal is coordinated via multiple neuromodulatory systems, such as norepinephrine^22^, acetylcholine^22^ and serotonin^30^, as well as shifts in local inhibitory and disinhibitory circuits^27,31,32^. Ethanol exposure affects both neuromodulation^18,33-35^ and inhibition in the brain^16,17,36-39^. Yet, the relationship between voluntary ethanol consumption and the coupling between cortical activity and arousal remains unexplored. Establishing this relationship is particularly important given the relevance of arousal states to stress and anxiety-like phenotypes^40-44^ as well as neuropsychiatric disorders like posttraumatic stress disorder (PTSD), which are well-established factors contributing to dysfunction of the PFC^45-47^ and AUD^41,48-50^.

Testing the effect of ethanol consumption on the activity of PFC neurons and their coupling to arousal requires simultaneous measurement of neuronal activity and arousal while animals voluntary consume ethanol. Several studies have utilized behavioral paradigms in which freely moving animals consume ethanol, allowing for sophisticated studies of neurophysiology during drinking, including extracellular electrophysiology^51-54^, one-photon mini-scopes^55^, fiber photometry^56^, and optogenetics^57,58^. To our knowledge, two-photon microscopy has not yet been leveraged to investigate changes in neurophysiology during volitional ethanol consumption. Two-photon microscopy complements these techniques by providing fine spatial and temporal resolutions for studying the effects of ethanol on the brain. It enables analysis of population-level neuronal activity at cellular resolution to study how ethanol consumption affects activity dynamics. It also enables longitudinal imaging of single cells and subcellular compartments, thereby providing information about the dynamics within single drinking sessions as well as over the longer time scales of days and weeks. However, a key drawback of two-photon microscopy is that it necessitates head-fixation for stable imaging, thereby limiting the scope of behaviors that can be studied.

We reasoned that mice with a history of binge-like drinking would transfer their behavior to drinking ethanol while being head-fixed under a two-photon microscope. We report here such a paradigm to open the possibility of applying two-photon microscopy and other recording modalities requiring head-fixation to the study of voluntary consumption of ethanol. In addition to two-photon imaging, we also collected pupil dynamics and licking behavior to correlate changes in neurophysiology in the PFC with changes in arousal and quantitative metrics of ethanol drinking. We targeted the anterior cingulate cortex (ACC) subdivision of the PFC for recordings. In humans, the ACC is affected by moderate doses of ethanol^12^ and its structural features and functional connectivity are prospectively associated with future drinking^59,60^. Moreover, the ACC is a key component of the cortical circuitry for interoceptive processing^61^, making it a prime candidate for assessing how coupling between arousal and cortical activity relates to voluntary ethanol behaviors. We find that mice consume as much or more ethanol during head-fixed drinking as they do during the drinking in the dark (DID)^1,2^ paradigm in their home cages. Furthermore, we find unexpected changes in neuronal activity as animals consume ethanol, especially with regards to its coupling to pupil dynamics indicative of changing arousal states. These data represent the first reported two-photon imaging of PFC networks while animals voluntarily consume ethanol as a head-fixed extension of the DID paradigm.

## Results

### Novel paradigm for studying voluntary ethanol consumption behavior in head-fixed mice

We designed a novel behavioral paradigm for head-fixed mice to enable two-photon calcium imaging during voluntary self-administration of ethanol. Mice were implanted with a head-fixation bar and chronic imaging window to provide optical access for two photon microscopy in the anterior cingulate cortex (ACC) subdivision of the prefrontal cortex. After recovery from surgery, mice were first allowed to self-administer ethanol using the ‘drinking in the dark’ (DID) paradigm^1,2^. Mice were provided access to a bottle containing 20% ethanol in drinking water (v/v) three hours into the dark phase of their light cycle for three hours every day (Fig. 1A). Mice gradually increased their ethanol intake during the DID (Fig. 1B). After eight days of DID, we habituated mice to head-fixation in daily sessions of 30 minutes for five days, during which mice continued to undergo DID. After habituation, the DID procedure was discontinued and mice instead self-administered ethanol while head-fixed under a two-photon microscope. The head-fixed drinking (HFD) paradigm was performed three hours into dark phase of their light cycle, at the same circadian time as the DID procedure.

**Figure 1.**
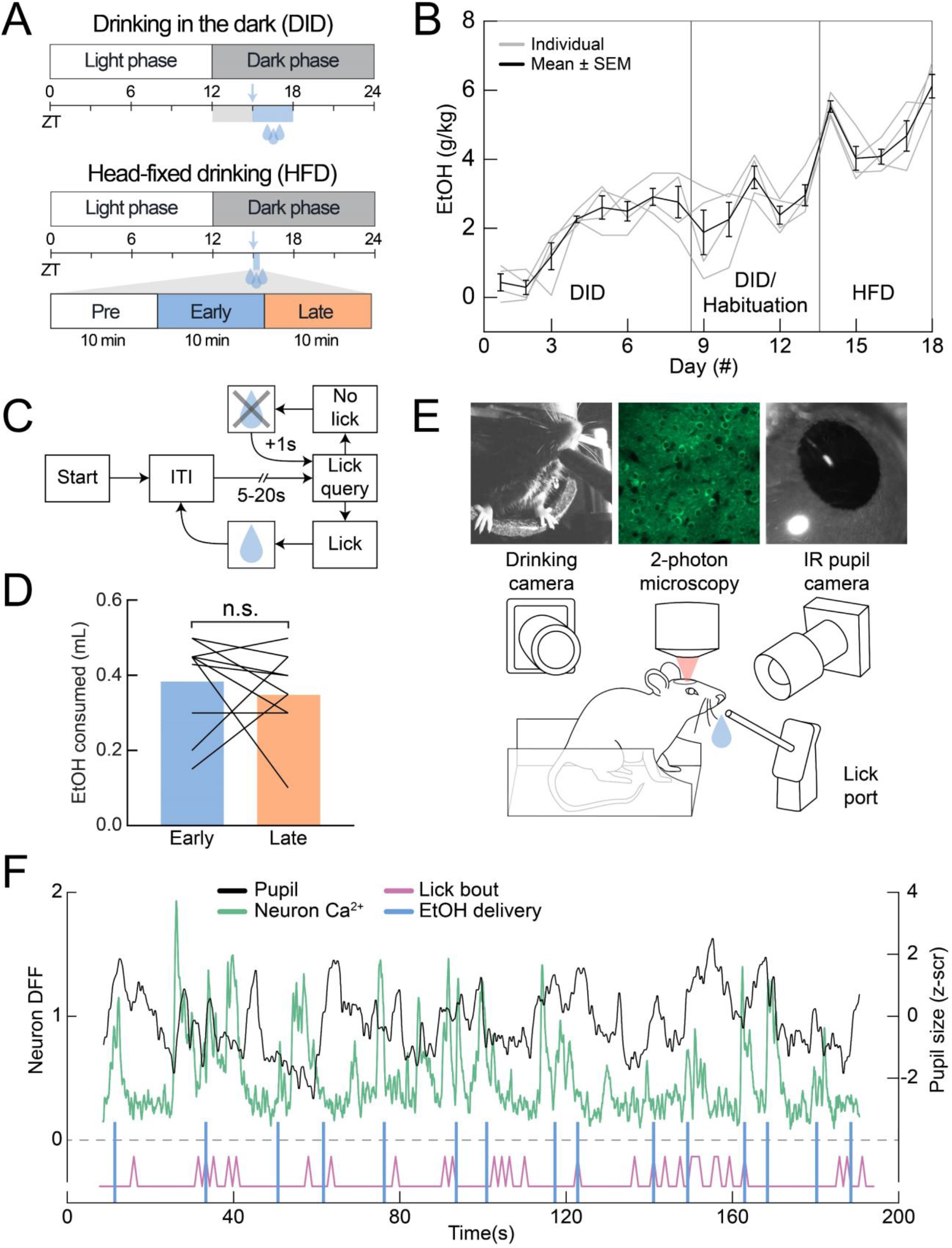
Head-fixed mice voluntarily consume ethanol during two-photon imaging. (A) Top: timeline of the ‘drinking in the dark’ paradigm (DID) in relation to zeitgeber time (ZT). Regular bottles were replaced with 20% ethanol (v/v) in drinking water for 3 hours beginning 3 hours into the dark phase (ZT15, blue) with habituation occurring prior (ZT12, gray). Bottom: timeline of head-fixed drinking (HFD). Animals were provided access at ZT15 for 3×10 minute sessions consisting of pre, early (blue), and late (orange) ethanol access. (B) Ethanol consumption across DID, DID with head-fixed habituation, and HFD periods. n = 4 mice. (C) Task structure for voluntary ethanol delivery (ITI, intertrial interval). (D) Volume of ethanol consumed during early and late drinking blocks (paired t-test, t(11) = 0.77, p = 0.46; n = 12 sessions from 4 mice). (E) Schematic of imaging setup (bottom) with examples of drinking camera, 2-photon imaging, and pupil camera.. (F) Example simultaneously measured DFF traces of an example neuron (green), pupil size (black), ethanol delivery (blue), and the observed lick bouts (magenta). n.s., not significant.

Head-fixed mice could collect drops of a 20% ethanol solution by licking a metal spout. We custom designed an electronic circuit to control ethanol delivery via a solenoid valve and to measure lick responses with a capacitive sensor (Fig. S1). This circuit interfaced with custom software, allowing us to control the timing and amount of ethanol delivered. Head-fixed drinking sessions were organized into trials, the structure of which was not cued to the animal (Fig. 1C). Each trial started with a randomized delay sampled from an exponential distribution (10s mean with cutoffs at 5s and 20s). Mice triggered a drop of ethanol by licking the spout during the query period (last 1s of the delay). Ethanol delivery was delayed by 1s until a lick was made during the query period, thereby requiring the animal to volitionally initiate ethanol delivery via licking. The un-cued trial structure and randomized delays introduced uncertainty in the ethanol delivery time, potentially incentivizing licking behavior. Thus, with this self-paced contingency, mice could acquire varying amounts of ethanol based on their level of engagement with drinking. Practically, it also ensured that mice collected the previously dispensed drop through licking before additional ethanol was delivered.

Mice underwent five days of HFD while being simultaneously imaged using two-photon microscopy. The imaging sessions were divided into three consecutive blocks consisting of a 10 minute pre-drinking baseline period followed by two 10 minute drinking blocks, which we refer to as early and late drinking, respectively (Fig 1A). Mice readily adapted to drinking while head-fixed and consumed large amounts of ethanol in the short span of 20 minutes (4.89±0.04 g/kg, n=20 sessions, 5 days, 4 mice; Fig. 1B; Supplementary Video 1). This volume corresponded to a binge-like level of blood-ethanol content (BEC; 199.3±62.19 mg/dL, n=4 mice, final day of imaging)^62^. We assessed whether mice front loaded ethanol consumption in this paradigm by comparing the amount consumed during the early and late drinking blocks. Across sessions, mice drank a similar amount of ethanol in each of the two drinking blocks (early: 0.39±0.03mL; late: 0.35±0.03mL; Fig. 1D). As mice voluntarily consumed ethanol, we measured the calcium responses of layer 2/3 excitatory neurons of transgenic mice (CaMKII-Cre x Ai148D) stably expressing the genetically encoded calcium sensor GCaMP6f (Fig. 1E, F). In addition, we used high-speed videography to measure changes in the pupil diameter as a proxy for momentary fluctuations in arousal^26,63,64^. Together with the high-resolution readout of licking behavior, this HFD paradigm allowed us to assess the effect of voluntary ethanol consumption on ACC activity as well as its coupling to simultaneously measured changes in arousal (Fig. 1F).

### Effect of ethanol consumption on single neuron and network level activity

Previous work suggests that AUD is associated with disrupted PFC processing and that ethanol consumption dysregulates PFC function^65^. We determined how voluntary ethanol consumption in this paradigm affects neuronal activity in the ACC. While the high spatial resolution of two-photon microscopy allowed us to repeatedly find the same field of views (FOVs) and longitudinally track the activity of the same neurons across days of drinking (Fig. S1), in this study we prioritized analysis of large populations of neurons and included responses from all recorded neurons. To determine the effect of ethanol consumption on ACC activity, we detected transient increases in GCaMP6f traces to quantify the frequency and amplitude of calcium events in individual neurons (Fig. 2A). Inspecting the activity of single neurons across the slow time scale of the entire imaging session showed diverse types of responses with drinking; while some neurons increased or decreased activity, others were unaffected (Fig. 2A). To assess how the ACC population as a whole responds to ethanol consumption, we compared the calcium event frequency of all recorded neurons during pre-drinking and late drinking. Overall, event frequencies were largely similar across these conditions (Fig. 2B). This lack of a population level effect is likely driven by heterogeneity in the response profiles of individual neurons, as suggested by the single neuron examples (Fig. 2A). The lack of a trial structure for the pre-drinking block makes it difficult to statistically assess changes in activity levels for individual neurons in an unbiased way. We addressed this issue by performing for each neuron a shuffle test which compared the observed difference in activity during pre-drinking and the two drinking blocks to the difference expected by chance given the overall activity in the entire recording period (see Methods). We found that ethanol consumption significantly modulated the activity of a subset of neurons, increasing the activity of 11.6±1.4% and decreasing the activity of 13.4±1.8% of neurons (p = 0.54, Wilcoxon signed-rank test; n = 12 sessions from 4 mice; Fig 2C-F).

**Figure 2.**
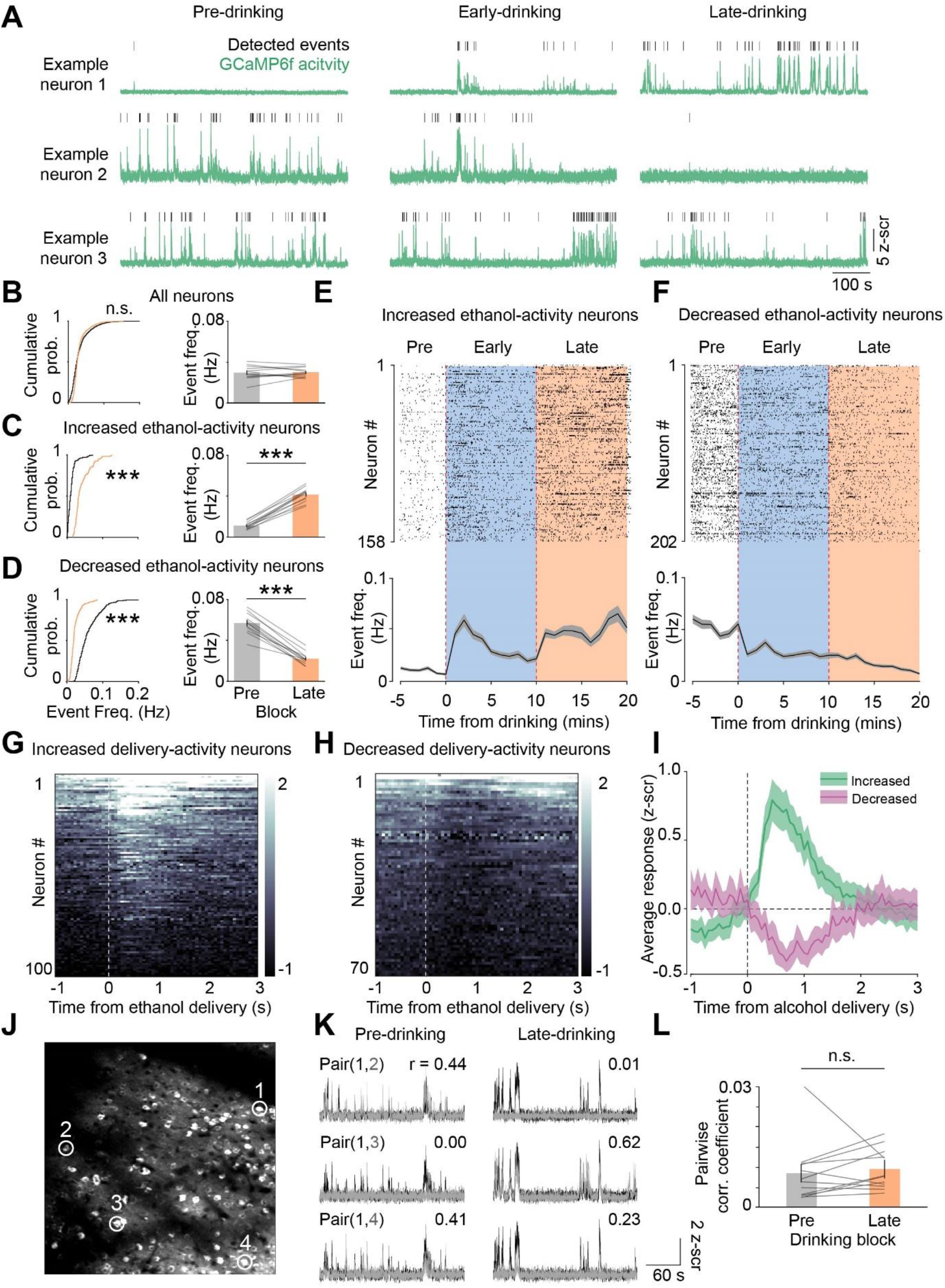
Effect of ethanol consumption on single neuron ACC activity and pairwise correlations. (A) Activity of three example neurons during pre, early, and late drinking. Detected calcium events (black) are shown overlaid on the z-scored DFF signal (green). (B) DFF event rates during pre (gray) and late (orange) drinking for all neurons. Cumulative probability (prob.) distribution (p = 9×10^−18^, k = 0.1638; n = 1471 neurons from 4 mice) and session-wide mean of events rates (p = 0.88, z = -0.16; n = 12 sessions from 4 mice) is shown. (C) Same as B, except for neurons with increased activity modulated by ethanol (prob. distribution, p = 10^−47^, k = 0.82, n = 158 neurons from 4 mice; mean event rates, p = 0.002; z = -3.06; n = 12 sessions from 4 mice). (D) Same as B, except for neurons with decreased activity modulated by ethanol (prob. distrubtion, p = 10^−43^, k = 0.69, n = 202 neurons from 4 mice; mean event rates, p = 0.002; z = 3.06; n = 12 sessions from 4 mice). (E) Raster plot showing detected events for neurons with significantly increased activity across drinking (top). Rows show activity for individual neurons. Mean event rates in 1 min bins averaged across all increased activity neurons are shown below. (F) Same as E, except for decreased activity neurons. (G) Color plot showing the trial-averaged activity of neurons (rows) showing increased activity around the time of ethanol delivery. (H) Same as G, except for neurons showing decreased activity around ethanol delivery. (I) z-scored DFF averaged across neurons with increased (green) and decreased (magenta) activity that are shown in G and H, respectively. (J) Example field of view (FOV), noting the spatial location of four representative neurons (white circles/numbers). (K) The activity of pairs of neurons shown in J for pre-drinking and late drinking blocks. (L) Mean pairwise correlation coefficients averaged for all unique pairs during individual pre- and late drinking blocks (p = 0.18, z = -1.33, n = 12 sessions from 4 mice; Wilcoxon signed-rank test). In B-D, comparison of cumulative probability distributions with two-tailed, two-sample Kolmogorov-Smirnov test and comparison of means with two-tailed Wilcoxon signed-rank test. ***p < 0.005, n.s., not significant.

The above analysis considered activity changes during drinking on the timescale of minutes, possibly reflecting the slow and cumulative effect of ethanol consumption throughout the session. Next, we addressed how activity is modulated on the faster timescale of individual ethanol delivery events. We aligned neuronal responses to the time of ethanol delivery and identified modulated neurons by comparing responses before and after delivery. Similar proportions of neurons had increased and decreased activity (6.6±3.8% & 4.3±3.5%, p > 0.05; Fig. 2G-I). Since mice licked the metal tube to collect the ethanol drop, this activity could be related to motor responses. If this were the case, we would expect that these same neurons respond to licking in general. Aligning neuronal responses to the onset of licking bouts around ethanol delivery (Fig. S2A) or licking during the inter-trial interval (Fig. S2B) did not show a consistent modulation in activity. Hence, the observed activity modulation likely reflects the delivery and/or receipt of ethanol. Together, these analyses show that ethanol delivery bidirectionally modulates ACC activity both at the slow time scale of minutes and at the sub-second time scale around ethanol delivery.

### Effect of ethanol consumption on pairwise neuronal correlations

Information processing in cortical networks is critically shaped by inter-neuronal correlations between pairs of neurons^66,67^. We took advantage of the large number of simultaneously recorded neurons in our dataset to test the effect of ethanol consumption on pairwise correlations. For each recording session, we computed the Pearson correlation coefficient between the activity of unique pairs of neurons (Fig. 2J, K). Visualizing the activity of single example pairs showed diverse changes, with correlations increasing, decreasing, or being unaffected during late drinking (last 5 minutes of the second drinking block) as compared to pre-drinking (Fig. 2K). To examine this process at the population level, we compared correlations averaged over all pairs recorded simultaneously in single behavioral sessions. Ethanol consumption did not significantly affect pairwise correlations (Fig. 2L). While pairwise correlations decreased as a function of distance between recorded pairs of neurons, there was no difference in this relationship between pre- and late drinking (Fig. S3A, B). Similarly, accounting for the average level of activity between neuron pairs did not show any differences (Fig. S3C). The percentage of pairs with significant correlations was similar between pre- and late drinking (Fig. S3D). Together, these analyses show that neuronal correlations in the ACC are largely unaffected by ethanol consumption when considering all recorded ACC neurons.

### Coupling between ACC activity and arousal during pre-drinking is prospectively associated with ethanol engagement

Our results thus far show that ethanol delivery modulates ACC activity. The activity of cortical neurons is profoundly affected by arousal^26,27^, which is reflected in changes in pupil size. We tested whether arousal-mediated modulation of ACC activity is related to ethanol consumption. Previous work shows that while activity in neurons of the posterior cortex (e.g., visual cortex) shows predominantly positive correlations with the pupil (i.e., increased activity during pupil dilations), frontal cortex neurons exhibit both positive and negative correlations (i.e., increased and decreased activity during pupil dilations, respectively)^27^. This suggests that subsets of frontal cortical neurons are activated as others are inhibited by fluctuations in arousal. Correlating neuronal calcium traces with pupil diameter during pre-drinking identified neurons with significant pupil-activity correlations, with individual neurons exhibiting positive, negative, or no correlation with changes in pupil diameter (Fig. 3A,B). Across the population, the pre-drinking activity of a similar proportion of neurons was positively and negatively correlated with changes in pupil diameter, albeit with large session-to-session variability (Fig 3C). The ability to detect activity-pupil correlations could depend on the level of neuronal activity, especially for neurons with negative correlations as this requires high levels of spontaneous activity when the pupil is constricted. We quantified the frequency and amplitude of detected calcium events and compared these measures between neurons with positive and negative correlations to pupil size. Overall, neurons correlated with pupil had similar level of activity during the pre-drinking period (Fig. 3D). Thus, subsets of ACC neurons have sufficient level of spontaneous activity to measure their positive and negative coupling to arousal.

**Figure 3.**
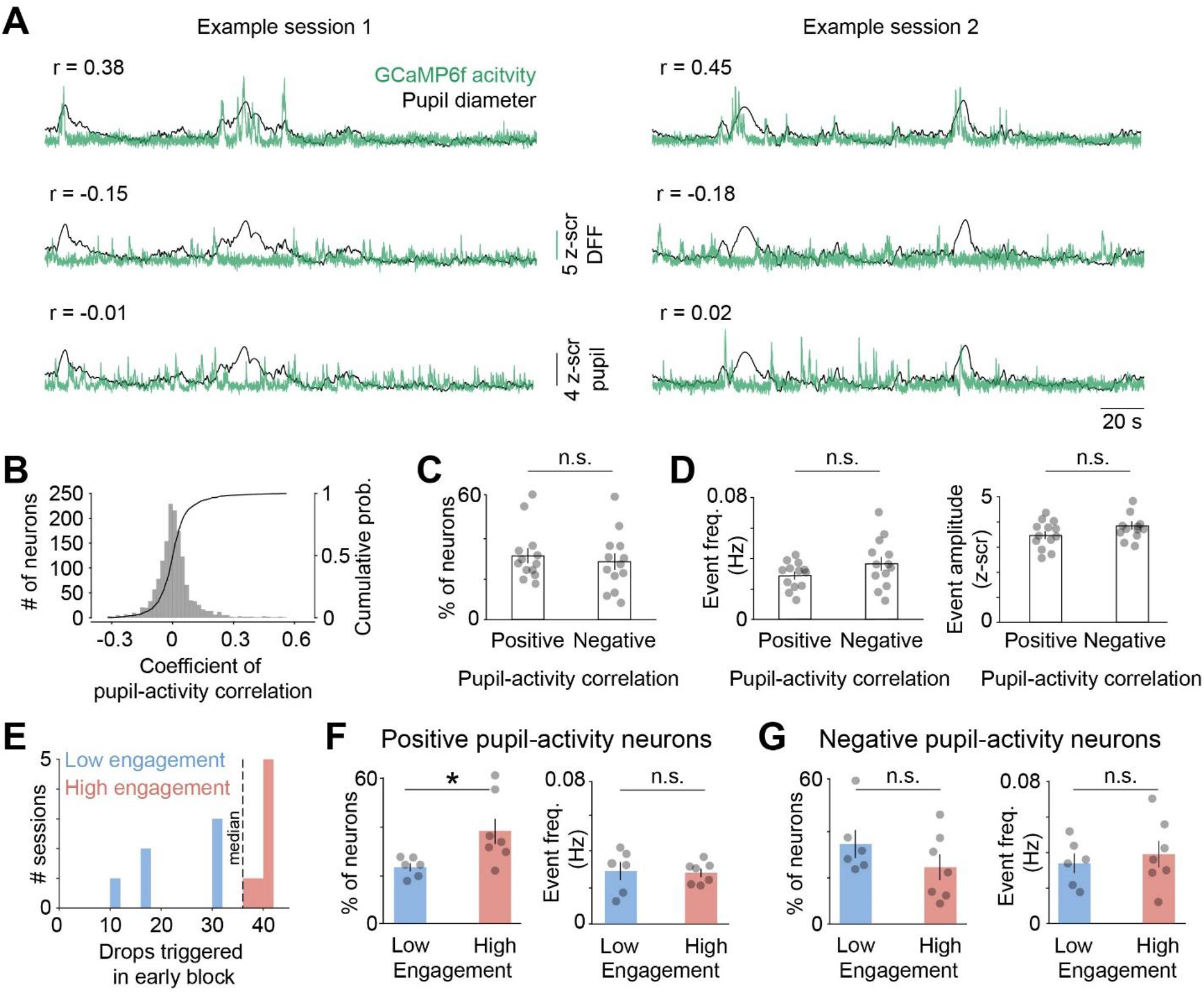
Pupil-activity correlation during pre-drinking is associated with subsequent engagement with ethanol. (A) Traces of pupil diameter (z-scored, black) overlaid on GCaMP6f DFF responses (z-scored, green) of representative neurons from two example sessions showing varying levels of pupil-activity coupling. (B) Histogram (left y-axis) and cumulative probability distribution (right y-axis) for Pearson correlation coefficient between fluctuations in neuronal activity and pupil diameter (n = 1519 neurons from 4 mice). (C) Percentage of neurons with significant positive and negative activity-pupil correlations (n = 13 sessions from 4 mice; p = 0.78, z = 0.28). (D) DFF event frequency (left; p = 0.20, z = -1.28) and amplitude (right; p = 0.20, z = -1.28) for positively and negatively coupled neurons (n = 13 sessions from 4 mice). (E) High (red) and low (blue) ethanol engagement sessions were defined based on median split of the number of ethanol drops triggered during early drinking (high, 7 sessions from 4 mice; low, 6 sessions from 3 mice). (F) Percentage of neurons with positive coupling to pupil in low and high engagement sessions (p = 0.02, z = -2.36) and their DFF event rates in the two sessions (p = 0.52, z = 0.64) (G) Same as F, but for neurons with negative pupil-activity correlations (percent of neurons, p = 0.23, z = 1.21; event rate, p = 0.83, z = -0.21). Statistical testing with two-tailed Wilcoxon rank-sum tests in all panels. *p < 0.05, n.s., not significant.

We addressed the relationship between cortical coupling to arousal during pre-drinking and the level of engagement with ethanol in the subsequent drinking block. In our paradigm, mice engage with ethanol by licking during the query period to trigger delivery. We defined low and high engagement sessions based on the median split of triggered ethanol delivery (Fig. 3E). The percentage of neurons with significant positive pupil-activity correlations during pre-drinking was higher in sessions preceding high levels of ethanol engagement as compared to sessions with low engagement (Fig. 3F). In contrast, the percentage of neurons with negative activity-pupil correlations was similar for low and high engagement sessions (Fig. 3G). There was no difference in the frequency of pre-drinking calcium activity preceding high and low sessions (Fig. 3F,G), suggesting that the overall level of activity in neurons is not associated with the level of ethanol engagement.

We further examined this phenomenon by aligning neuronal activity to the time of individual pupil dilation events. To perform this analysis, we first identified individual pupil events (Fig. 4A). Importantly, there were no differences in the frequency, amplitude, or duration of pupil events during pre-drinking blocks preceding low and high engagement sessions (Fig. 4B). This suggests that differences in pre-drinking pupil dynamics do not account for the level of subsequent ethanol engagement. As expected, neurons with positive pupil-activity correlations were activated as the pupil started to dilate, reaching maximal activity before the peak of pupil dilation; meanwhile, neurons negatively coupled to arousal had decreased activity during pupil dilations (Fig. 4C). Importantly, neurons with positive pupil-activity correlations had increased pupil-aligned responses preceding high as compared to low engagement sessions (Fig. 4C, D). In contrast, the pupil aligned activity of negatively correlated neurons was similar for both sessions (Fig. 4C, E).

**Figure 4.**
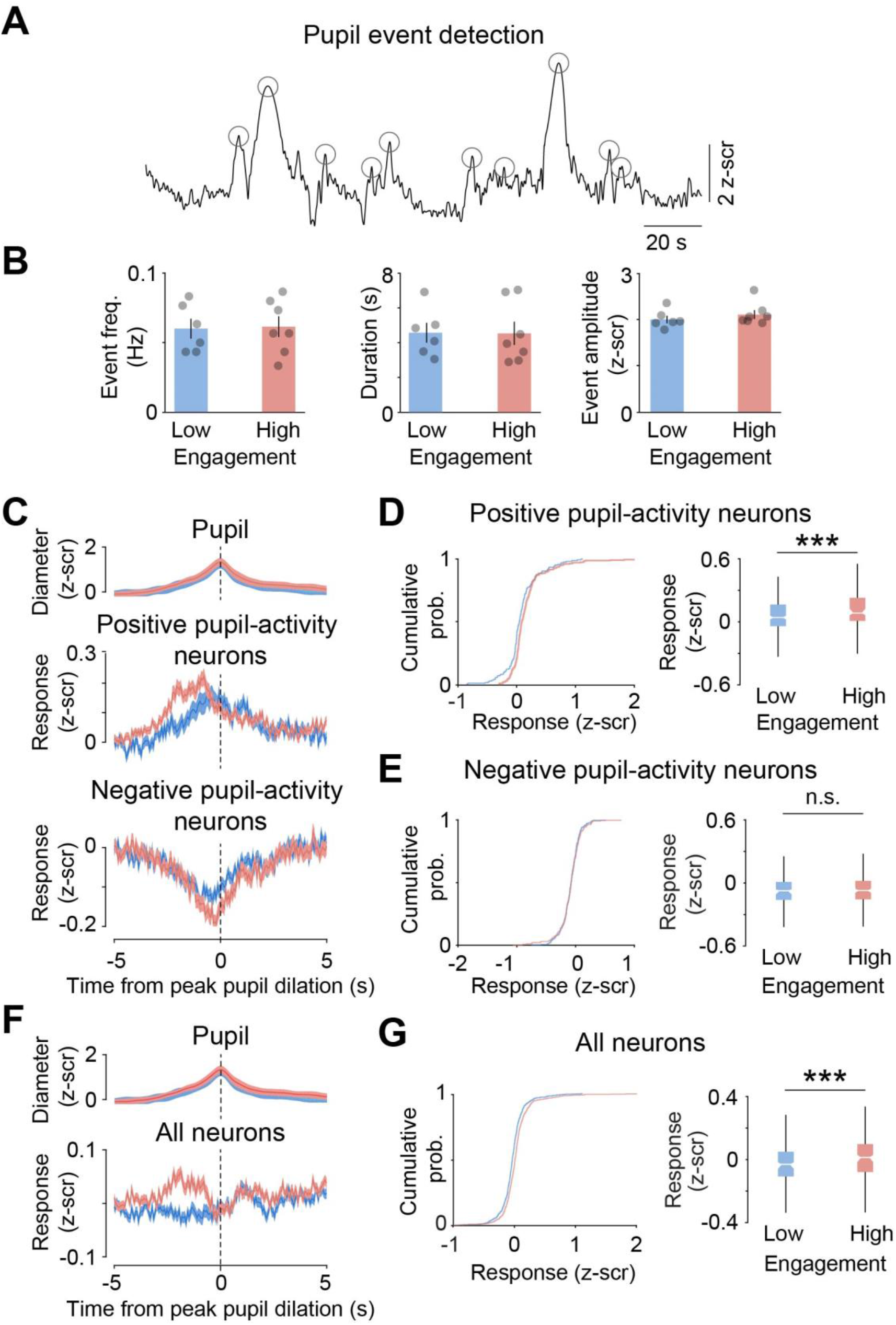
Pupil-aligned neuronal responses during pre-drinking are associated with subsequent ethanol engagement. (A) Trace of pupil diameter (z-scored, black) from an example session showing peaks of individual dilation events (circles). (B) Quantification of pupil event rate (p = 0.94, z = -0.07), duration (p = 0.62, z = 0.5), and amplitude (p = 0.35, z = -0.92) for low (blue) and high (red) engagement sessions (n = 7 high sessions from 4 mice, 6 low sessions from 3 mice). (C) Session-averaged pupil events (top) and neuronal responses from positively (middle; low, n = 171 neurons from 3 mice; high, n = 297 neurons from 4 mice) and negatively (bottom; low, n = 242 neurons from 3 mice; high, n = 212 neurons from 4 mice) coupled neurons, with signals aligned to the peak of pupil dilation events. Responses from low (blue) and high (red) engagement sessions are shown. (D) Cumulative probability distribution (left panel; p = 0.00013, k = 0.21) and median (right panel; p = 0.002, z = -3.17) pupil-aligned responses averaged over a window of -2 to -1s relative to peak pupil dilation. (E) Same as D, except for negatively coupled neurons (cumulative distribution: p = 0.92, k = 0.05; box plot: p = 0.85, z = -0.19). (F) Same as C, except for all neurons regardless of correlation with the pupil (low, n = 718 neurons from 3 mice; high, n = 801 neurons from 4 mice). (G) Same as D, except for all neurons (cumulative distribution, p = 2×10^−9^, k = 0.16; box plot, p = 5×10^−9^, z = -5.84). Box plot elements: center line, median; box limits, upper and lower quartiles; whiskers, 1.5x interquartile range; outliers not shown). Comparisons in B and the box plots in D, E, G with two-tailed Wilcoxon rank-sum test; significance testing of cumulative distributions in D, E, G with two-tailed, two-sample Kolmogorov-Smirnov test. ***p < 0.005, n.s., not significant.

Thus far, we have focused on neurons that show significant pupil-activity correlations. We next assessed overall neuronal-arousal coupling in the ACC population by analyzing the pupil aligned responses of all neurons. As expected, combining neurons with both positive and negative pupil correlations as well as uncoupled neurons showed less arousal modulation than observed before (Fig 4F). Despite this, activity around pupil dilation events was higher during pre-drinking blocks preceding high engagement sessions as compared to low sessions (Fig. 4G). Together, we find that the level of positive coupling between ACC activity and pupil-linked arousal during pre-drinking is prospectively associated with subsequent engagement with ethanol. Importantly, this association is not explained by differences in pre-drinking pupil dynamics (Fig. 4B). Rather, it reflects the effect of arousal on cortical circuits, predominantly on neurons that are activated by arousal (Fig. 4C, D, F, G).

### Ethanol consumption reconfigures the coupling between arousal and neuronal activity

The above results establish neuronal-arousal coupling as an important factor for ethanol related behavior in this paradigm. We determined how ethanol consumption affects this coupling. Our previous analysis (Figs. 3, 4) only examined neuronal-arousal coupling during the pre-drinking period, leaving open the possibility that additional neurons in the population become coupled to arousal with drinking. Hence, we identified arousal modulated neurons as ones with significant pupil-activity correlations either during pre-drinking or the subsequent drinking blocks. Across the population, a similar proportion of neurons had positive or negative coupling to arousal (positive: 55±5.2%; negative: 45.5±5.6%; p = 0.14, two-sided Wilcoxon rank-sum test; n = 12 behavioral sessions from 4 mice). To track the evolution of neuronal-arousal coupling during the entire imaging session, we computed pupil-activity correlation coefficients in five-minute bins across drinking (Fig 5A). For cells with positive coupling, drinking had no significant effect on pupil-activity correlations (Fig. 5A, B). In contrast, drinking decreased negative coupling to arousal (Fig. 5D, E). Importantly, the frequency of calcium events for both positively and negatively coupled neurons were unaffected by drinking (Fig. 5C, F), suggesting that the observed changes in coupling to arousal are not due to gross changes in neuronal activity.

**Figure 5.**
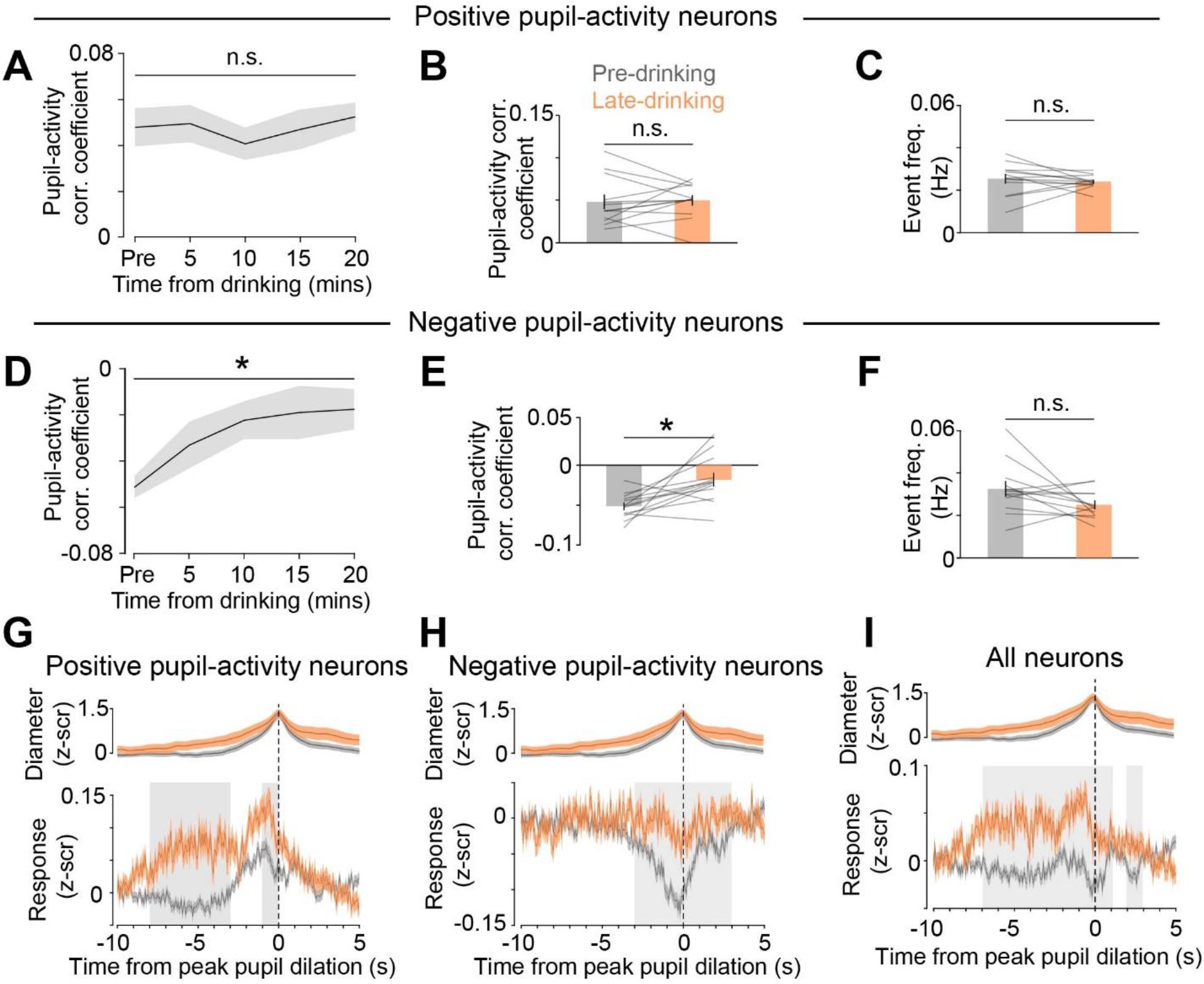
Voluntary ethanol consumption disrupts neuronal-arousal coupling. (A) Session-averaged correlation (corr.) coefficients for positive pupil-activity modulated neurons, quantified in 5 min bins across drinking (H(4,55) = 2.13, p = 0.71; n = 12 sessions from 4 mice). (B) Average pupil-activity correlation coefficients during pre-drinking and last 5 minutes of late-drinking (p = 0.81, z = -0.24). (C) DFF event frequency for pre-drinking (gray) and late (orange) drinking block (p = 0.48, z = 0.71). (D, E, F) Same as A, B, C except for negative pupil-activity neurons (panel D, H(4,55) = 9.7, p = 0.046; panel E, p = 0.012, z = -2.51; panel F, p = 0.14, z = 1.49). (G) Pupil responses (z-scored, top) and pupil-aligned activity (z-scored, bottom) of neurons with significant positive pupil-activity correlations during pre- and late drinking. Activity was compared in 1s windows using two-tailed Wilcoxon signed-rank test; gray shading denotes windows with significant difference between the two blocks using Bonferroni corrected α-value of 0.0033 (0.05/15). (H, I) Same as G, except for negative pupil-activity neurons (H) and all recorded neurons (I). Significant testing with Krusal-Wallis test in A and D, and with Wilcoxon signed-rank test in B, C, E, and F. *p < 0.05, n.s., not significant.

We investigated which factors contribute to the observed changes in pupil-activity correlations with ethanol consumption. Drinking had moderate effects on the pupil dynamics, leading to a non-significant decrease in the pupil event frequency and increase in event duration (Fig. S4). A change in the frequency of pupil fluctuations without a concomitant change in the frequency of neuronal activity could in principle account for the observed decrease in pupil-activity correlations. We controlled for this by analyzing pupil-aligned neuronal responses. The pupil-aligned responses of positively coupled neurons were increased in late drinking compared to pre-drinking, largely mirroring the broadening of pupil dilation events observed with drinking (Fig. 5G). In contrast, the activity of negatively coupled neurons exhibited less arousal-mediated modulation with drinking, showing a reduction in decreased activity observed around pupil dilation events compared to pre-drinking (Fig. 5H). These results suggest that ethanol consumption shifts the balance of neuronal-arousal responses to favor excitatory coupling. In agreement, pupil-aligned activity of all ACC neurons analyzed without regard to their coupling to arousal showed increased activity with drinking as compared to pre-drinking (Fig. 5I). Together, these results demonstrate that ethanol consumption asymmetrically reconfigures the coupling between arousal and neuronal activity, weakening negative and strengthening positive neuronal-arousal coupling.

## Discussion

We developed a novel behavioral paradigm for voluntary ethanol consumption in head-fixed mice (Fig. 1). This allowed the application of two-photon microscopy to measure diverse facets of neuronal signaling in the ACC, components of which were prospectively associated with ethanol engagement in addition to being affected by drinking. Freely moving mice readily transitioned from the DID paradigm to consuming ethanol while head-fixed (Fig. 1A-C). Measuring neuronal activity in layer 2/3 excitatory neurons of the ACC showed that ethanol self-administration modulates neuronal activity, both at the sub-second time scale during consumption of individual drops of ethanol and at the slower time scale of minutes throughout drinking (Fig. 2). Interestingly, there was a bidirectional relationship between ethanol consumption and neuronal-arousal coupling. Increased neuronal excitation in response to arousal during pre-drinking was prospectively associated with higher levels of ethanol engagement (Figs. 3, 4). Importantly, pupil dynamics were similar preceding high and low ethanol engagement sessions; hence, this coupling effect is due to cortical responses to arousal rather than the level of arousal itself. In parallel, drinking shifted the balance of neuronal-arousal coupling, weakening inhibitory and strengthening excitatory responses to arousal (Fig. 5). Although small subsets of ACC neurons did show activity modulation with drinking, the observed changes in neuronal-arousal coupling were not accompanied by changes in the overall activity level of neurons. This demonstrates that for most ACC neurons, ethanol consumption largely exerts a selective effect on responses to momentary fluctuations in internal state shifts rather than a non-specific effect on their activity. Together, the fact that ethanol increased excitatory responses to arousal and that increased responses to arousal before drinking are prospectively associated with high ethanol engagement suggests ethanol-mediated reconfiguration of neuronal-arousal coupling as an important signature of neuroadaptations relevant for AUD.

The extension of the DID paradigm to head-fixed drinking allows the application of two-photon microscopy to the study of voluntary ethanol-related behaviors. The high spatial resolution of two-photon imaging allows longitudinal tracking of physiological adaptations simultaneously with ethanol engagement (Fig. S1B). While longitudinal experiments with cellular-resolution have been performed with one-photon microendoscopy in freely moving animals^55^, the GRIN lenses required for this technique have a limited depth of imaging compared to two-photon imaging through chronic windows. Moreover, a key advantage of using head-fixation is the ability to collect precise behavioral measures of ethanol consumption including simultaneous high-speed acquisition of pupil dynamics, locomotion, lick microstructure, whisking, and limb kinetics^68,69^. Combination of two-photon imaging with specific transgenic mouse lines^70,71^ and viral vectors for projection-specific labeling^72^, along with the expanding gamut of optical sensors for measuring neuronal activity as well as extracellular release of neurotransmitters^73^, will allow future experiments to study the effect of voluntary ethanol consumption on processing by cell-specific circuits. Indeed, two-photon imaging was recently used to examine changes in cortical visual processing in anesthetized mice that had previously undergone chronic exposure to ethanol^74^. Similarly, two-photon calcium imaging is beginning to identify how systemically injected ethanol modulates neuromodulator release and non-neuronal cell types^75^.

The head-fixation required for two-photon imaging poses limitations. Recent work shows that head-fixation in mice initially increases levels of the stress-related hormone corticosterone, which normalizes over days with habituation^76^. We previously measured circulating corticosterone levels under head fixation relative to other stressors. While habituated head-fixation elicits significantly less stress hormone release compared to restraint stress paradigms, it is still significantly elevated from baseline in naïve mice^77^. Another limitation is the temporal resolution of measured neuronal activity using two-photon microscopy. Calcium-based neuronal activity profiles are orders of magnitude slower than electrophysiological techniques, though our head-fixed approach can be used with emerging tools for voltage imaging or high-density electrophysiology such as NeuroPixel probes^78^ that also require head-fixation. Given these limitations, two-photon microscopy is a complementary approach to electrophysiological studies in freely-moving paradigms with specific unique advantages.

While we used the head-fixed drinking paradigm to study various aspects of activity in the ACC and volitional ethanol consumption, we envision our approach as a baseline head-fixed paradigm that can be easily modified to address other important questions relevant for AUD, including stress-ethanol interactions^79^ and instrumental ethanol seeking^80^. We found that mice consumed large quantities of ethanol during head-fixed drinking. Given the well-appreciated relationship between stress and excessive drinking in humans and in animal models^48,81,82^, one possibility is that the mild stress associated with head-fixation promotes drinking in this paradigm. Hence, this paradigm, combined with other stressors, can be leveraged to dissect the relationship between ethanol consumption and stress as well as the neuronal-arousal coupling described here. This paradigm may also facilitate the study of compulsive ethanol-related behaviors^51,83,84^ in head-fixed mice. Here, drops of ethanol were available after random delays drawn from an exponential distribution. Such delays have a flat subjective hazard rate and disallow subjects from predicting when an event will occur^85,86^. In this regard, our paradigm is similar to conditioning experiments using random interval reinforcement schedules, which are thought to promote compulsive behavior^87^. This suggests that the high levels of drinking observed in our model may be extended to investigate punishment-resistant compulsive drinking, at least in a subset of mice. Future modifications that allow testing this and other AUD-related behaviors in head-fixed mice will allow application of two-photon microscopy to these questions.

We found that voluntary ethanol consumption in awake mice both increased and decreased the activity of subpopulations of ACC neurons at the slow time scale of minutes as well as on the faster time scale of individual ethanol delivery events (Fig. 2). In anesthetized rats, intraperitoneal injection of ethanol dose-dependently inhibited the activity of PFC neurons recorded via extracellular electrophysiology^88^. Subsequent work directly compared the effect of ethanol injections in anesthetized vs. awake conditions and found decreased activity during anesthesia but generally no overall changes in activity levels during wakefulness^89^. This suggests that ethanol has a particularly depressive effect on PFC activity in the anesthetized state. Compared to systemic ethanol injections, voluntary ethanol consumption may induce more heterogenous activity patterns that potentially reflect multiple processes including the intent to drink, olfaction, gustatory/consummatory responses, olfaction, and others. Future adaptations in the behavioral paradigm that carefully distinguish between these processes are needed to clarify the precise nature of this observed activity.

We focused on the activity of layer 2/3 excitatory neurons, however the observed coupling of neuronal activity to the pupil suggests involvement of inhibition. Processing in cortical circuits is crucially coordinated by precise spatiotemporal inhibition, with recent studies highlighting the role of molecularly defined inhibitory neurons in neuronal-arousal coupling^70,90,91^. Vasointestinal peptide (VIP) expressing interneurons provide inhibition onto other inhibitory neurons, including the somatostatin (SOM) expressing subtype^32,92-94^. Activation of pyramidal neurons during arousal is thought to reflect a disinhibitory microcircuit motif wherein arousal-mediated activation of VIP cells causes inhibition of SOM cells and disinhibition of pyramidal neurons, thereby providing a potential mechanism for positive responses to arousal. In the frontal cortex, direct VIP inhibitory inputs onto pyramidal cells are proposed to mediate the negative responses to arousal^27^. Thus, one possibility is that ethanol exposure reconfigures neuronal-arousal coupling by reshaping these inhibitory circuits and shifting the balance between inhibition and disinhibition of pyramidal neurons. Alterations in inhibition are commonly found in response to ethanol exposure, with the general finding that inhibition is reduced^95,96^. Importantly, recent work has started addressing how specific subtypes of cortical interneurons are affected, uncovering changes in SOM neuron intrinsic excitability and synaptic outputs that could be particularly relevant for neuronal-arousal coupling^97,98^.

In addition to altered inhibition, cortical brain state shifts are strongly correlated with brain-wide release of neuromodulators including norepinephrine, acetylcholine, corticosterone, and serotonin^22,24,28,30^. Numerous studies have implicated dysregulated neuromodulatory activity with AUDs, suggesting that neuronal coupling to arousal may be a signature of neuromodulation indicative of drinking behaviors^33,49,99^. Relatedly, acute exposure to ethanol profoundly disrupts the activity of locus coeruleus noradrenergic neurons^75,100^, which form a key component of the brain’s neuromodulatory arousal system^101^. Future work will be needed to address the causal relationship between neuromodulatory systems, arousal-coupled cortical activity, and ethanol consumption.

Although considerable mechanistic research has been done on the acute cellular effects of ethanol, including effects on the fast time scales of ion channel activity and synaptic transmission^102-107^, as well as the chronic long-term circuit neuroadaptations^36,108-110^, relatively few studies have studied changes in the intervening time course of cortical information processing. Our work bridges these timescales by focusing on how acute ethanol engagement modifies circuit dynamics on the order of internal state shifts, which are highly relevant for behavioral functions often associated with the PFC. Better understanding the effect of ethanol on this intermediate timescale will further clarify the relationship between the myriad of behavioral and neuronal adaptations that comprise AUD.

## Figures

**Supplementary Figure 1.**
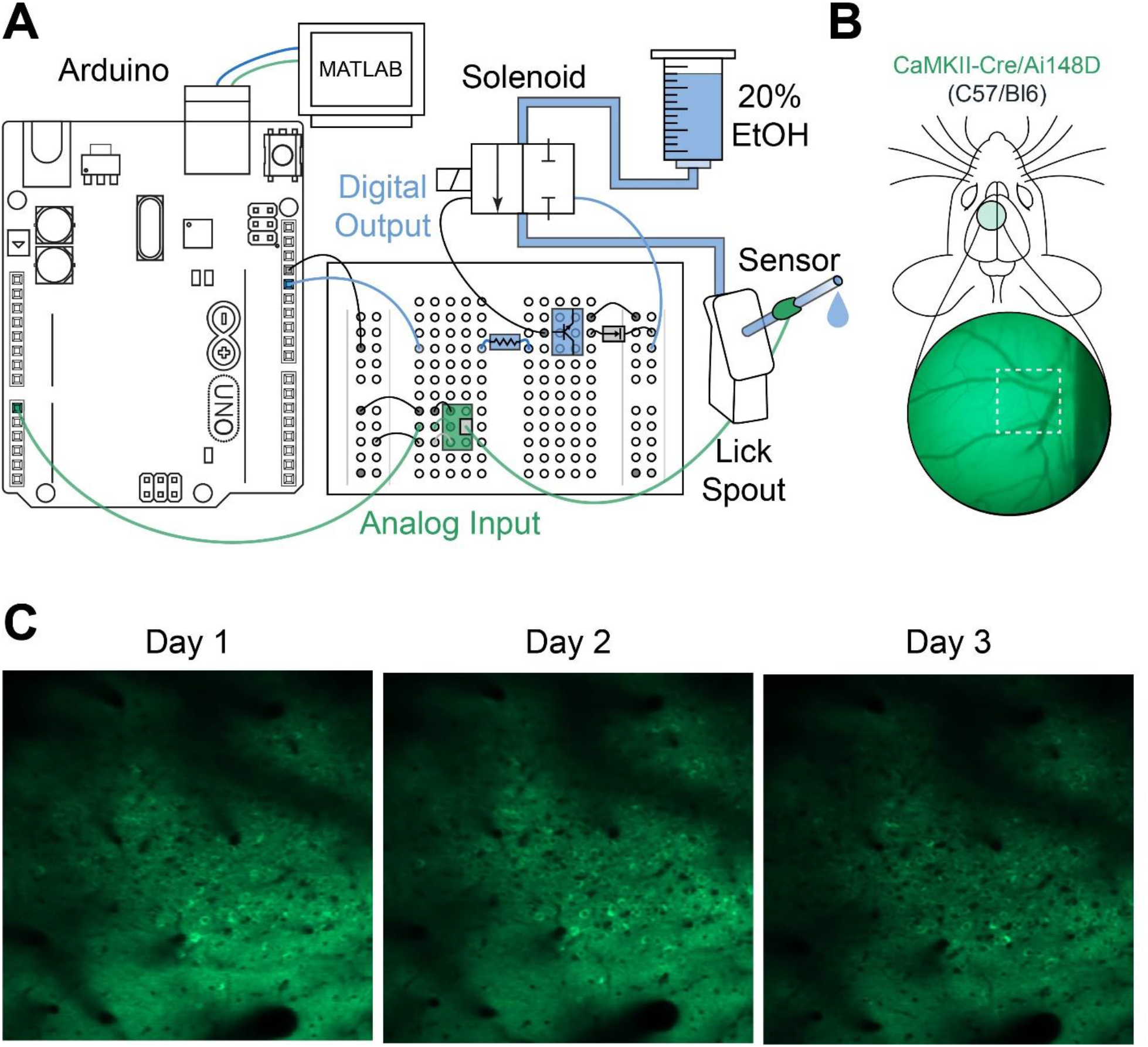
Voluntary ethanol engagement with cortical imaging over days. (A) Circuit diagram for lick detection (analog input, green) and ethanol delivery (digital output, blue). (B) Example low-magnification field of view of anterior cingulate cortex (ACC) and imaging region (white dashed box). (C) Example imaging of excitatory ACC neurons across 3 days. Coordinates of 0.5mm anterior to Bregma and 0.3mm lateral to the midline were targeted using recorded imaging depth, vasculature shadow patterns, and background fluorescence of neuronal somata.

**Supplementary Figure 2.**
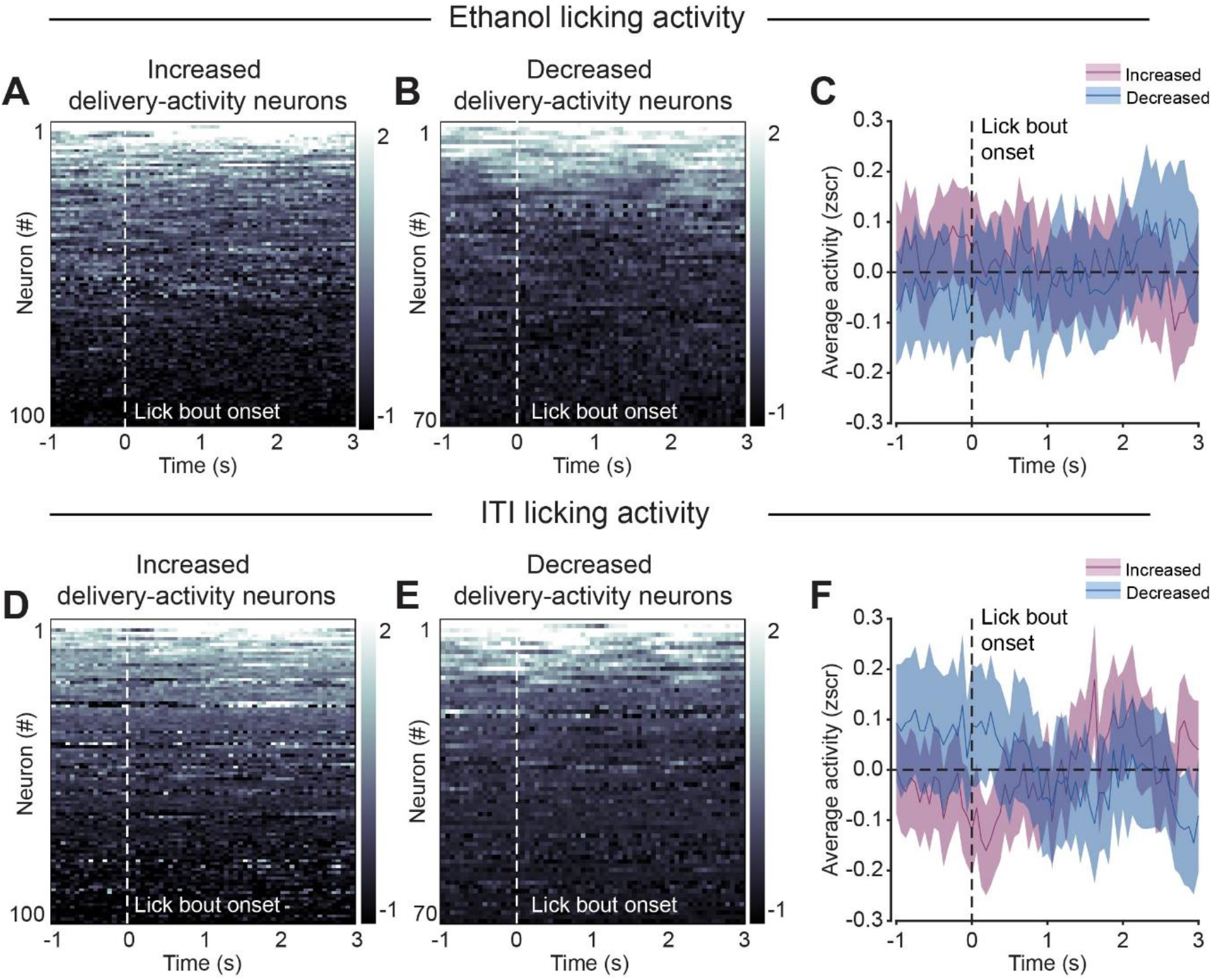
Licking does not significantly modulate neuronal activity. (A) Activity of neurons with increased ethanol delivery-activity shown in Fig. 2G but aligned to licking bouts occurring around ethanol delivery. (B) Same as A, but for neurons with decreased delivery-activity neurons shown in Fig. 2H. (C) Averaged DFF responses of increased (green) and decreased (magenta) delivery-activity neurons (as in Fig. 2I), but aligned to lick bouts around ethanol delivery. (D, E, F) Same as A, B, C but aligned to licking bouts in the inter-trial interval (ITI).

**Supplementary Figure 3.**
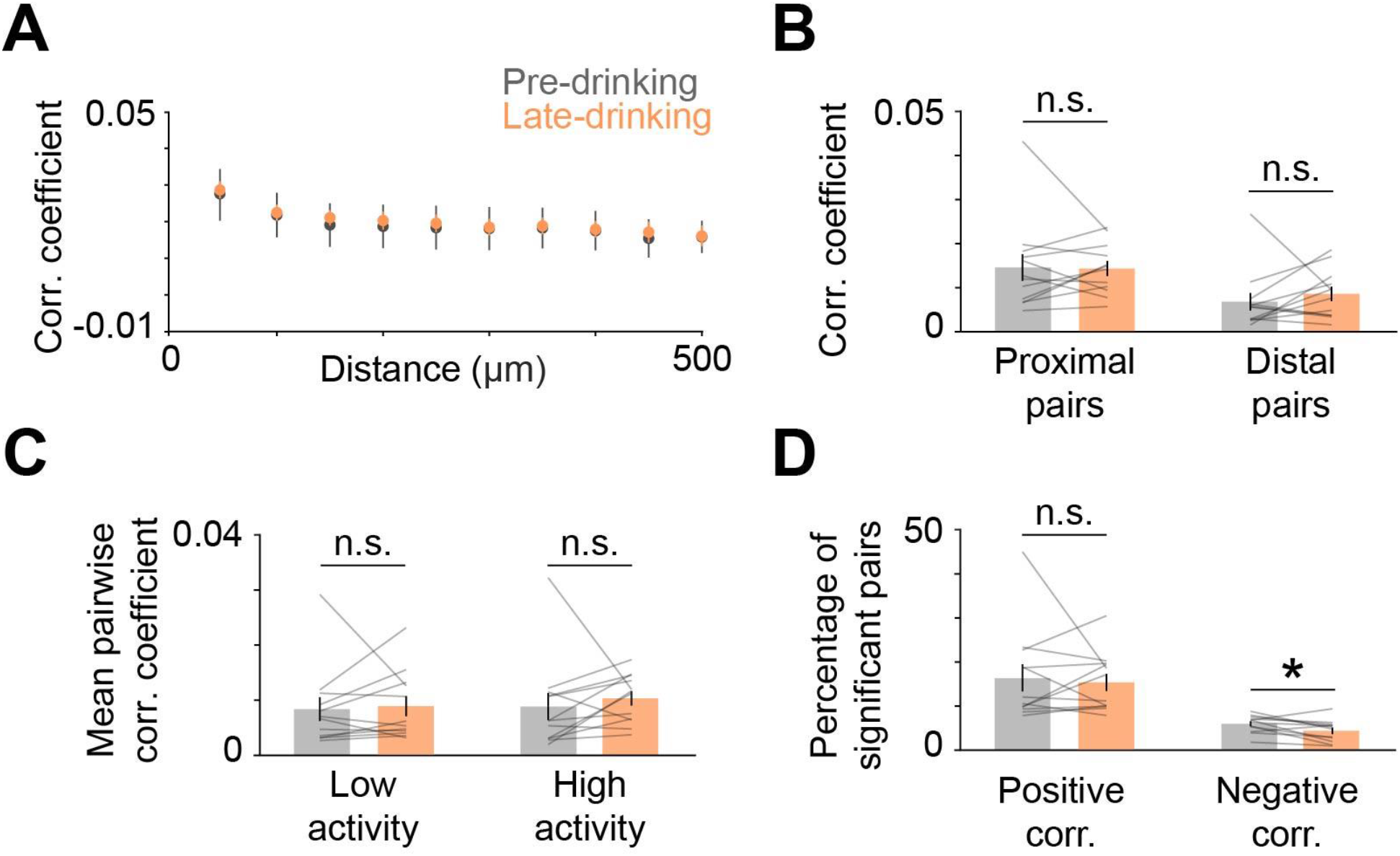
Effect of ethanol consumption on pairwise activity correlations. (A) Session-averaged pairwise correlation coefficients between all unique pairs of simultaneously recorded neurons binned by the distance between neurons in each pair (bin size, 50µm). (B) Activity correlations of proximal (<50µm apart) and distal (>300µm apart) pairs during pre (gray) and late (orange) drinking (proximal, p = 0.58, z = -0.55; distal, p = 0.21, z = -1.26). (C) Comparison of correlation coefficients of pairs with low and high activity, defined based on the median split of the calcium event frequency averaged across the neurons in a pair (low activity, p = 0.88, z = -0.15; high activity, p = 0.07, z = -1.80). (D) Percentage of neuronal pairs with significant positive or negative pairwise correlations (positive, p = 0.94, z = -0.08; negative, p =.0.041, z = 2.04). Error bars are standard error of the mean (n = 12 sessions from 4 mice). Significant testing with Wilcoxon rank-sum test; *p < 0.05; n.s., not significant.

**Supplementary Figure 4.**
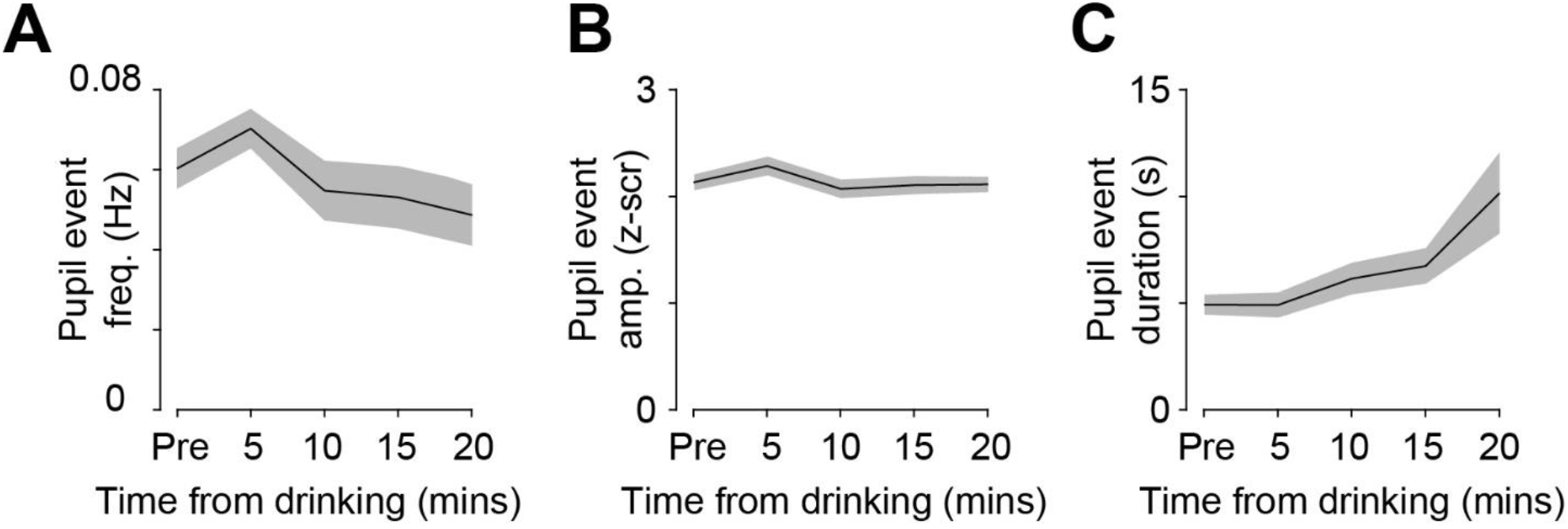
The effect of drinking on pupil dynamics. (A-C) Frequency (H(4,55) = 6.82, p = 0.14), amplitude (H(4,55) = 4.71, p = 0.22), and duration (H(4,55) = 8.95, p = 0.06) of detected pupil dilation events in 5 min bins across drinking (n = 12 sessions from 4 mice). Significance testing with one-way Kruskal-Wallis test.

**Supplementary Video 1. Home-cage video of mice after voluntary head-fixed ethanol consumption**.

## Methods

### Mice

Animals expressing GCaMP6f under the CaMKII promoter were generated by crossing the commercially available Camk2a-Cre (005359, Jackson) and Ai148D (030328, Jackson) mouse lines, both on a C57/Bl6 background. Animals were maintained in their home cages under a 12/12 reverse light/dark cycle with *ad libitum* access to standard mouse chow and water. All animals used in this study were ∼P60 male mice. All animal procedures were performed in strict accordance with protocols approved by the MIT Division of Comparative Medicine and conformed to NIH standards.

### Surgical Procedures

Surgeries were performed under isoflurane anesthesia (3% induction, 1.5% maintenance) and body temperature was maintained at 37.5°C using a temperature controller (ATC2000, World Precision Instruments). Animals were dosed with slow-release buprenorphine (0.1mg/kg) prior to surgery, and meloxicam (1mg/kg) every 24 hours post-surgery for 72 hours or until fully recovered. Once stably anesthetized, animals were head-fixed in a stereotaxic frame (51500D, Stoelting), scalp hair removed, and scalp sterilized using alternating betadine scrub and 70% ethanol solutions. A portion of the scalp was removed and conjunctive tissue cleared after treatment with hydrogen peroxide. A 3mm diameter craniotomy was then drilled over centered over left ACC/M2 (from bregma, AP: 1.0mm, ML: 1.0mm). A chronic cranial window was then implanted that consisted of a 5mm diameter coverslip glued to two 3mm coverslips (Warner Instruments) using optical UV-cured adhesive (61, Norland). The window was carefully lowered with the 5mm coverslip on top and firmly held in the craniotomy using the stereotax while adhered to the skull using dental acrylic mixed with black ink (Metabond, Parkell). Once the dental acrylic had cured around the cranial window, a custom stainless-steel head-fixation plate was mounted around the cranial window and cured with dental acrylic. Animals were then allowed to recover in their own cage with a warm water blanket and moistened food chow. Mice were singly housed for the remainder of the experiment and continued to recover for one week before beginning the drinking in the dark paradigm.

### Drinking in the dark (DID) paradigm

Following recovery from window implantation, mice were introduced to the drinking in the dark paradigm (DID) as previously described for 13 days. Mice were weighed, and their water bottles replaced with 15ml conical tubes containing 20% ethanol (v/v) diluted in the mouse drinking water 3 hours into their dark phase (ZT15). Tubes were filled with ∼10mL of ethanol solution, fitted with a custom rubber stopper and standard lick spout, and then weighed before being placed in each cage. One drop of ethanol solution was allowed to drip to ensure the displacement of air. A control cage without any mice was also fitted with an ethanol drinking tube to account for the displacement drop and evaporation. Mice were then left alone for 3 hours during which they could voluntarily consume the ethanol solution. After 3 hours, the ethanol tubes were removed, weighed, and regular drinking water bottles returned. The amount of ethanol consumed was calculated by subtracting the final weight from the initial weight of each tube to get a session difference. The control tube difference was then subtracted from each mouse tube difference, and then consumption computed as ethanol consumed (g) per weight of the animal (kg). After 9 days of normal DID exposure, mice were head-fixed on the two-photon imaging rig for 30 minutes (ZT 12-15) prior to the DID paradigm (ZT15) for 5 days to habituate the mice to the imaging setup.

### Head-fixed drinking paradigm

After animals completed 13 days of DID, last 5 days of which they were habituated to head-fixation, mice were imaged while voluntarily consuming ethanol under the two-photon microscope. Animals were head-fixed on an elevated platform with a lickspout delivering 20% EtOH (v/v) in mouse drinking water positioned within easy access for licking. The lickspout was made from a brass tube (3.97mm diameter, 8128, K&S Precision Metals) that was wrapped with conductive wire and connected to a capacitive sensor (P1374, Adafruit) integrated to a breadboard and connected to an Arduino board (Uno Rev3, A000066, Arduino) as an analog input and recorded via custom MATLAB scripts. Ethanol delivery was initiated by custom MATLAB scripts that sent a digital signal via the Arduino to toggle a transistor on the breadboard (IRF520PBF, Digi-Key) and open a 12V solenoid (VAC-100 PSIG, Parker). Ethanol solution was maintained in a graduated syringe, gravity fed into the solenoid, and calibrated to deliver a small drop (∼8µL) with each trigger. Following each session, total ethanol delivered was compared between the graduated syringe and the calculated trigger volume to ensure accurate quantification of total ethanol delivered (for detailed circuit diagram and session structure see Fig. 1 and Supp. Fig. 1).

For the initial sessions after switching from DID to head-fixed drinking, mice could lick the spout to receive ethanol soon after head-fixation. To determine how drinking affects ACC activity relative to before drinking, in later sessions we introduced a 10 minute pre-drinking imaging block in which animals were allowed to lick the spout but no ethanol was delivered. Following the pre-drinking block, animals underwent two subsequent 10 minute drinking blocks in which licking the spout could deliver a drop of ethanol. The drinking blocks were structured such that after initiation, there was a pseudorandom delay before a lick query (exponential distribution with a 10s mean and cut-offs at 5s and 20s). If the animal had licked within 1s prior to the query, a drop of ethanol solution was dispensed, otherwise an additional delay of 1s was imposed before another lick query. Hence, the animal had to lick during the query period to trigger ethanol delivery. This design promoted continuous licking of the spout in order to obtain more ethanol, thereby limiting unconsumed ethanol delivery. Moreover, the self-paced design allowed us to assess the animal’s level of engagement with ethanol by quantifying the number of drops that were triggered. A camera (LifeCam, Q2F-00013, Microsoft) was positioned to record licking behavior and to validate that ethanol delivery was consumed. To quantify changes in arousal, an infrared CMOS camera (DCC1545M, ThorLabs) was fitted with a telecentric lens (SilverTL, 58-430, Edmund Optics) and positioned over the pupil for high-speed acquisition (20Hz) via custom MATLAB scripts. The pupil was illuminated by an infrared LED array (LIU780A, ThorLabs). Ethanol delivery, image acquisition, and pupil imaging were all synchronously triggered at the start of the session to accurately align temporal epochs across measurements.

We analyzed data from sessions with imaging during both pre-drinking and drinking blocks. We had 13 such sessions, which were included in the analysis of pre-drinking activity (Figs. 3, 4). 1 session had poor quality imaging data in the late drinking block and hence was excluded from data presented in Figs. 2 and 5.

### Measurement of blood ethanol concentration

Blood ethanol concentration (BEC) was quantified after the final imaging session. Immediately the session, animals were rapidly anesthetized by isoflurane, decapitated, and whole trunk blood collected in tubes lined with EDTA (BD Microtainer, 365974, Becton, Dickinson and Company) and placed on ice. Whole blood was then centrifuged at 3000xg for 10 minutes at 4°C. Separated plasma was then aliquoted and immediately stored at -80°C for further analysis. Quantification of BEC was performed using a colorimetric assay as previously described. Ethanol standards and plasma samples were diluted in sample reagent (all reagents obtained from Millipore Sigma): 100mM KH2PO4 (P3786), 100mM K2HPO4 (P3786), 0.7mM 4-aminoantropyrine (A4382), 1.7mM chromotropic acid (27150), 50mg/L EDTA (E4884), and 50mL/L Triton X100 (X100). Working reagent was created by mixing alcohol oxidase from *Pichia* (5kU/L, A2404) and horseradish peroxidase (3kU/L, 77332) with sample reagent and mixed with samples on a 96-well plate. Following 30 minutes of incubation at room temperature, the samples and standards were read on a standard plate reader (iMark, Bio-Rad Laboratories) at 595nm. Samples and standards were run 6x in parallel and BEC calculated according to the standard curve.

### Two-Photon calcium imaging

GCaMP6f fluorescence from neuronal somas was imaged through a 16x/0.8 NA objective (Nikon) using resonance-galvo scanning with a Prairie Ultima IV two-photon microscopy system. Image frames were collected as 4-frame averages at 480×240 pixel resolution an acquisition rate of 16Hz. Excitation light at 900nm was provided by a tunable Ti:Sapphire laser (Mai-Tai eHP, Spectra-Physics) with ∼10-20 mW of power at sample. Emitted light was filtered using a dichroic mirror (collected with GaAsP photomultiplier tubes (Hamamatsu). Layer 2/3 GCaMP6f-expressing neurons were imaged with 1.5x optical zoom, 120-200μm below the brain surface. Neuronal activity in the ACC was collected at the following coordinates: ∼0.5mm AP, ∼0.5mm ML.

### GaMP6f fluorescence signal processing

We used the software Suite2P^111^ for semi-automatic detection of neuronal somas from calcium imaging movies. Movies from the three imaging blocks were concatenated together and the non-rigid translation function in Suite2P was used to correct for x-y translations that may have occurred between blocks. Suite2P detects neuronal regions of interest (ROI) by clustering neighboring pixels with similar fluorescence time courses. Moreover, it provides for each detected neuron a neuropil mask which surrounds the detected ROI and excludes other detected neuronal ROIs. The automatically detected ROIs were manually curated using the GUI such that ROIs without clear visual evidence for neuronal somas were rejected and neurons missed by the algorithm were added manually.

To minimize the contribution of the neuropil signal to the somatic signal, corrected neuronal fluorescence at each time point t was estimated as F_t_ = Fraw_soma_t_ – (0.3 x Fraw_neuropil_t_)^112^. The DFF (ΔF/F) for each neuron was calculated as ΔF/F(t) = 100 x (F(t) – F_0_)/F_0_, where F_0_ represents the mode of the distribution of fluorescence values (estimated using the MATLAB function ‘ksdensity’). The resulting DFF trace was z-scored. We identified individual calcium events as transient increases in the z-scored DFF signal. Using the ‘findpeaks’ function in MATLAB, we detected events with minimum peak prominence of 2.5 z-scored DFF and minimum width of 3 imaging frames (∼200ms) at half-height of the event peak. All analyses either used the z-scored DFF or detected calcium event frequency and amplitude.

### Analysis of change in neuronal activity with ethanol consumption

We tested how ethanol consumption affects neuronal activity over the slow time scale of minutes across the entire imaging session. A challenge with this analysis is that the pre-drinking block has no trial structure, making it difficult to align activity to specific events and statistically compare how drinking affects activity. While one strategy is to compare inter-event intervals between pre-drinking and drinking blocks, we reasoned that the sparse cortical activity observed in individual blocks would be a limiting factor and produce false negatives. Hence, we instead addressed this issue by devising a shuffle test. This test circularly shifted traces of detected calcium events in time by a random amount in intervals of 30s, thus maintaining the temporal structure of activity while randomizing the timing at which it occurred. We reiterated this process 1000 times for each neuron. On every iteration, we computed the difference in event frequency between pre-drinking and 1) the first drinking block; and 2) the second drinking block. This allowed us to generate null distributions for the difference in event frequency expected by chance given the overall activity level of the neuron. The two-tailed p-value for each drinking block was computed as the proportion of activity changes in the null distribution that were as or more extreme than the experimentally observed change on either side of the distribution. Neurons with p < 0.05 for either drinking block were considered significant and classified as positively or negatively modulated depending on how their activity changed with drinking.

We also tested how individual ACC neurons were modulated on a faster time scale, around the time of ethanol consumption and licking. We aligned neuronal activity to the time of ethanol delivery and compared responses 1s before and after delivery using a one-tailed Wilcoxon signed-rank test to identify positively or negatively modulated neurons. Neurons with p < 0.01 were considered significant. We similarly aligned responses to licking bouts occurring at any time, during a 4s window after ethanol delivery, or during the inter-trial delay (i.e., 4s after ethanol delivery). Licking bouts were defined as two or more consecutive licks that occurred with a delay of less than 500ms.

### Pairwise neuronal correlation analysis

We assessed the effect of ethanol consumption on Pearson correlations between the z-scored DFF traces for all unique pairs of neurons in each recording session. To facilitate comparison between drinking and pre-drinking blocks, activity 3s after ethanol delivery in the drinking blocks was not included in the analysis. Correlations were computed using the last 5 minutes of the pre and late drinking blocks. Pair-wise correlations of all simultaneously recorded pairs in a single session were averaged together separately for the pre-drinking and drinking blocks. These session-averaged values were then compared between the blocks using a Wilcoxon signed-rank test. We quantified pair-wise correlations as a function of the distance between neurons in each pair. We calculated the Euclidean distance between each pair of neurons as the length of a straight line connecting the center coordinates of each neuronal ROI. We defined proximal pairs as neurons less than 50 µm apart and distal pairs as neurons with more than 300 µm distance between them. We also analyzed whether the overall level of activity in the pair of neurons is related to ethanol’s effect on pair-wise correlations. We did a median-split for the average event frequency for each pair and compared pair-wise correlations separately for pairs with high and low levels of activity. Lastly, we quantified and compared the proportion of neuron pairs with significant positive or negative pair-wise correlations during pre-drinking and post-drinking (p < 0.05).

### Neuronal-arousal coupling analysis

We performed two types of analyses to study the relationship between neuronal-arousal coupling and ethanol consumption: 1) correlation between fluctuations in pupil size and neuronal activity; 2) analysis of pupil-aligned neuronal activity. The pupil was imaged at a frequency 20Hz. We downsampled the pupil signal to match the frequency of two-photon calcium imaging data (∼16 Hz) using the MATLAB function ‘interp1’. For activity-pupil correlation analysis, we determined the Pearson correlation coefficient between neuronal activity and traces of z-scored pupil diameter. To evaluate the association of neuronal-arousal coupling during the pre-drinking block with subsequent drinking, we separately analyzed sessions with high and low levels of drinking, which were defined based on the median split of the total number of ethanol drops triggered by the animal in the first drinking block. Neurons were defined as positively or negatively modulated if they had a significant correlation with the pupil (p < 0.05) and depending on the sign of their correlation coefficient. Pre-drinking activity-pupil correlations were quantified for the second half of the pre-drinking block (i.e., last 5 minutes).

For tracking how correlations evolve across pre-drinking and drinking, we computed activity-pupil correlations coefficients in 5 min bins starting from the last 5 minutes of the pre-drinking to the end of the late drinking block. To facilitate comparison between pre-drinking and drinking blocks, neuronal and pupil data from 3s after each ethanol drop delivery during the drinking block were excluded from the analysis. Neurons with significant positive or negative correlations in any of the bins were included in the analysis (p < 0.05).

To analyze pupil-aligned neuronal activity, we first identified individual dilation events in traces of z-scored pupil diameter using the MATLAB function ‘interp1’ with a threshold prominence of 1 z-score. This allowed us to quantify the pupil dilation event rate, duration, and amplitude, in addition to identifying the time at which the peak of the dilation event occur. Neuronal responses were aligned to this peak time. For pre-drinking analysis, pupil-aligned responses for each neuron were averaged for all pupil events occurring in the last 5 mins of the block; for late drinking, last 5 mins of the second drinking block was used.

## Data availability

The data that support the findings of this study are available from the corresponding author upon reasonable request.

## Code availability

Custom code used in this work is available from the corresponding author upon reasonable request.

## Acknowledgements

This work was supported by grants from National Eye Institute F32EY028028 to G.O.S.; from National Institute of Mental Health R00MH104716 to E.V. and MH112855 to R.H.; and National Institute on Alcohol Abuse and Alcoholism U01AA025481 to E.V. We thank Mriganka Sur for generously providing the use of his laboratory for these experiments.

## Author contributions

G.O.S. and R.H. conceived the project, designed the experiments, and developed the concepts presented. G.O.S. performed the experiments with contributions from R.H. and E.V. R.H. performed the analysis with contributions from G.O.S., I.L. and M.N. E.V. provided feedback on overall experimental design and data interpretation. R.H. and G.O.S. wrote the manuscript with input from all authors.

